# 501Y.V2 and 501Y.V3 variants of SARS-CoV-2 lose binding to Bamlanivimab *in vitro*

**DOI:** 10.1101/2021.02.16.431305

**Authors:** Haolin Liu, Pengcheng Wei, Qianqian Zhang, Zhongzhou Chen, Katja Aviszus, Walter Downing, Shelley Peterson, Lyndon Reynoso, Gregory P. Downey, Stephen K. Frankel, John Kappler, Philippa Marrack, Gongyi Zhang

## Abstract

We generated several versions of the receptor binding domain (RBD) of the Spike protein with mutations existing within newly emerging variants from South Africa and Brazil. We found that the mutant RBD with K417N, E484K, and N501Y exchanges has higher binding affinity to the human receptor compared to the wildtype RBD. This mutated version of RBD also completely abolishes the binding to a therapeutic antibody, Bamlanivimab, *in vitro*.

## Main

The SARS-CoV-2 virus has infected over one hundred million people (COVID-19 patients) and caused more than two million deaths to date (WHO_COVID-19_patients 2020). The number of affected people continue to grow rapidly, emphasizing the need for rapid use of effective vaccines. Although two mRNA (Pfizer-BioNTech COVID-19 Vaccine and MODERNA respectively) Spike protein based vaccines have been approved for emergency use in the USA (Polack et al. 2020; Baden et al. 2020), the increasing number of Spike variants that have appeared around the world raise concerns about the continued efficacy of the vaccines (SARS-COV-2-Variants 2021). It has been reported that ~90% of broadly neutralizing anti-SARS-CoV-2 antibodies from COVID-19 patients engage the receptor binding domain (RBD) of the virus Spike protein (Piccoli et al. 2020). Monoclonal antibodies specifically targeting the native form of the Spike developed by Regeneron and Eli Lilly have been approved by the FDA for emergency use (Baum et al. 2020; Gottlieb et al. 2021; Bamlanivimab 2020; casirivimab and imdevimab 2020). An N501Y variant of SARS-CoV-2 (B.1.1.7, 20I/501Y.V1), first emerging in the United Kingdom and now spreading to the rest of the world recently, appears much more contagious than the original version (SARS-COV-2-Variants 2021). We found that this single mutation of N501Y confers an ~10 times fold increase of affinity between RBD and ACE2 (Liu et al. 2021). However, this mutation does not affect its binding to the therapeutic antibody, Bamlanivimab (Liu et al. 2021). We concluded that the increase of high binding affinity may account for the high infection rate of the United Kingdom variant while both vaccines and the therapeutic antibody Bamlanivimab still should remain their efficacy to combat this newly emerging variant. However, this same N501Y mutation is also found in a variant (B.1.351, 20H/501Y.V2) from South Africa and a variant (P1, 20J/501Y.V3) from Brazil with additional mutations within the RBD (SARS-COV-2-Variants 2021). Unfortunately, the two additional mutations, K417N and E484K, are also critical residues involved in the interactions between RBD and ACE2 as well as Bamlanivimab. It has been reported that a COVID-19 patient was infected a second time by the new variant from Brazil (Resende, Bezerra, and al. 2021). These information raise big concerns whether current vaccines and therapeutic antibodies still remain efficacy. Here, we address the basis of how these two mutations affect the binding of RBD to ACE2 and Bamlanivimab.

As we reported early that Y501Y-RBD derived from the United Kingdom variant has an ~10 times fold increased binding affinity (0.566nM) toward ACE2 compared to the wildtype (5.768 nM) (Liu et al. 2021). It was reported that both the South African variant and the Brazilian variant are also highly contagious as the United Kingdom variant (Tegally et al. 2020). We asked whether these two additional mutations within the RBD region effect the binding affinity between RBD and ACE2 and account for the high infection rate. On the basis of Y501-RBD, two additional mutations, K417N and E484K, were introduced and expressed in 293F cells as previously reported (Liu et al. 2021). Purified protein N417/K848/Y501-RBD was subjected to binding assays to ACE2 on a Biacore machine. Interestingly, the binding affinity (~2.935 nM) between N417/K848/Y501-RBD and ACE2 is much lower than that of Y501-RBD and ACE2 (0.566 nM) though still ~2 time folds higher than that between wildtype RBD (N501-RBD) and ACE2 (~5.768 nM) (**Fig. 1A**). Nevertheless, this ~2 fold binding affinity increase may partially account for the higher infectious rate presenting in South Africa and Brazil. It is of great interest on how these two additional mutations weaken the association between RBD and ACE2. To investigate the contribution of each additional mutation to the binding affinity, we carry out the following experiments. First, we introduce only E484K mutation into the existing N501Y-RBD to generate a double mutation of RBD (484K/501Y-RBD) and expressed it in 293F cells. Again, purified protein was subjected to Surface Plasmon Resonance binding assays on a Biacore machine to examine binding affinity between 484K/501Y-RBD and ACE2. As this E484K mutation brings a positively charged residue lysine to replace a negatively charged residue glutamic acid, we expected a drastic drop of binding affinity between 484K/501Y-RBD and ACE2. To our surprise, the binding affinity between this double mutation version of RBD (K484/Y501-RBD) and ACE2 is ~0.654 nM (**Fig. 1B**), an affinity that is similar to that of Y501-RBD and ACE2 at ~0.566 nM as previously reported. It suggests this mutation does not affect the binding between RBD and ACE2. To verify this conclusion, we explored the detailed interactions of K484/Y501-RBD and ACE2 and carried out molecular docking based on the published complex structure of RBD and ACE2 (Lan et al. 2020). Clearly enough, there could be a weak salt bridge (4.4Å, longer than 4.0 Å) formed between E484 of RBD and K38 from ACE2. After conversion of E484 to K484, this weak salt bridge is gone. It is likely that the slight affinity drop (from 0.566 nM to 0.674 nM) of K484/Y501-RBD is due to the loss of this weak interaction (**Fig 2A, 2B**). What happens if we introduce another mutation, K417N, on the basic 501Y-RBD? Similar to before, a double mutation version with K417N and N501Y protein (N417/Y501-RBD) was produced and subjected to binding to ACE2 on the Biacore machine.

**Figure 1.**
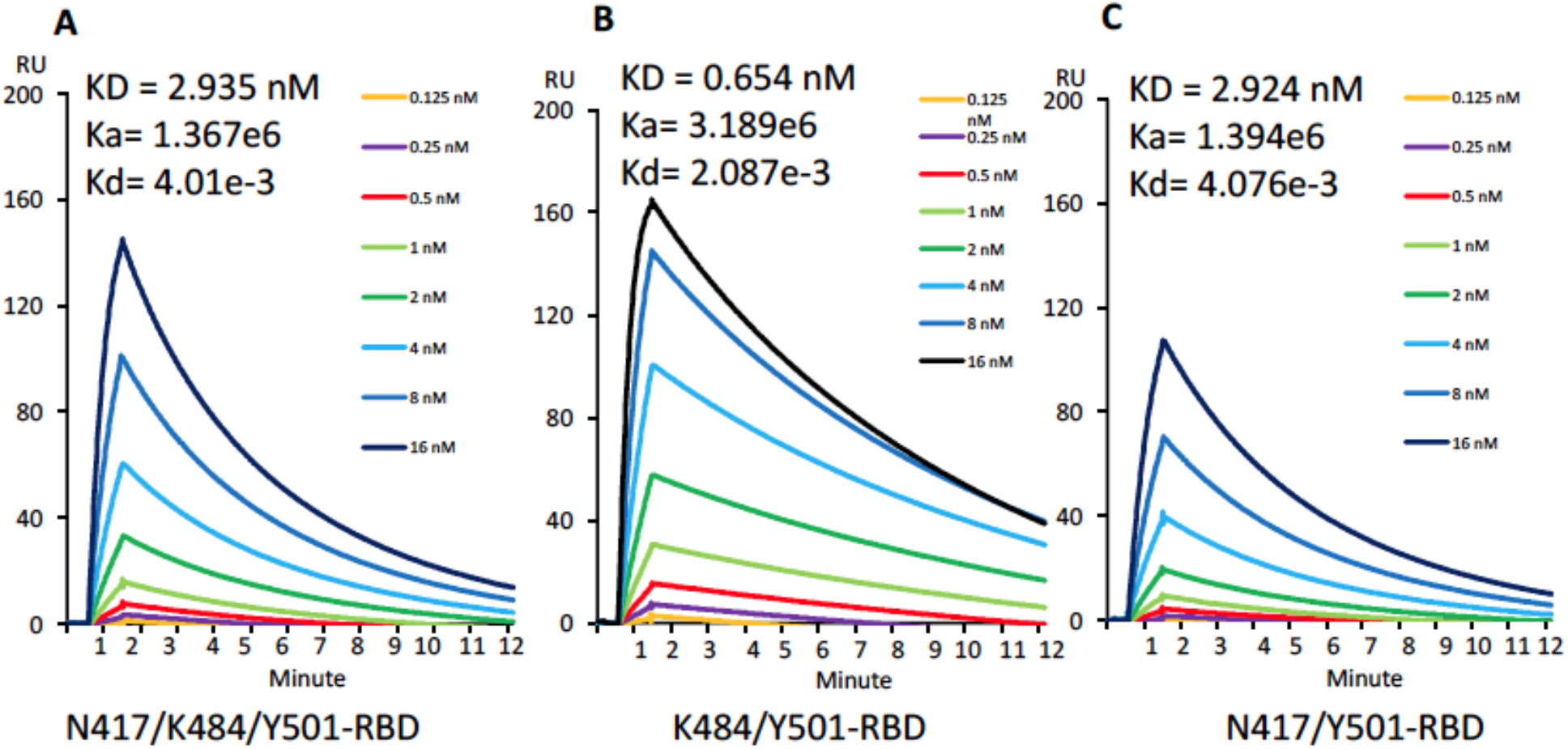
The basis of binding between mutated RBDs and ACE2. A. The Binding of N417/K484/Y501-RBD and ACE2. B. The binding of K484/Y501-RBD and ACE2. C. The binding of N417/Y501-RBD and ACE2.

**Figure 2.**
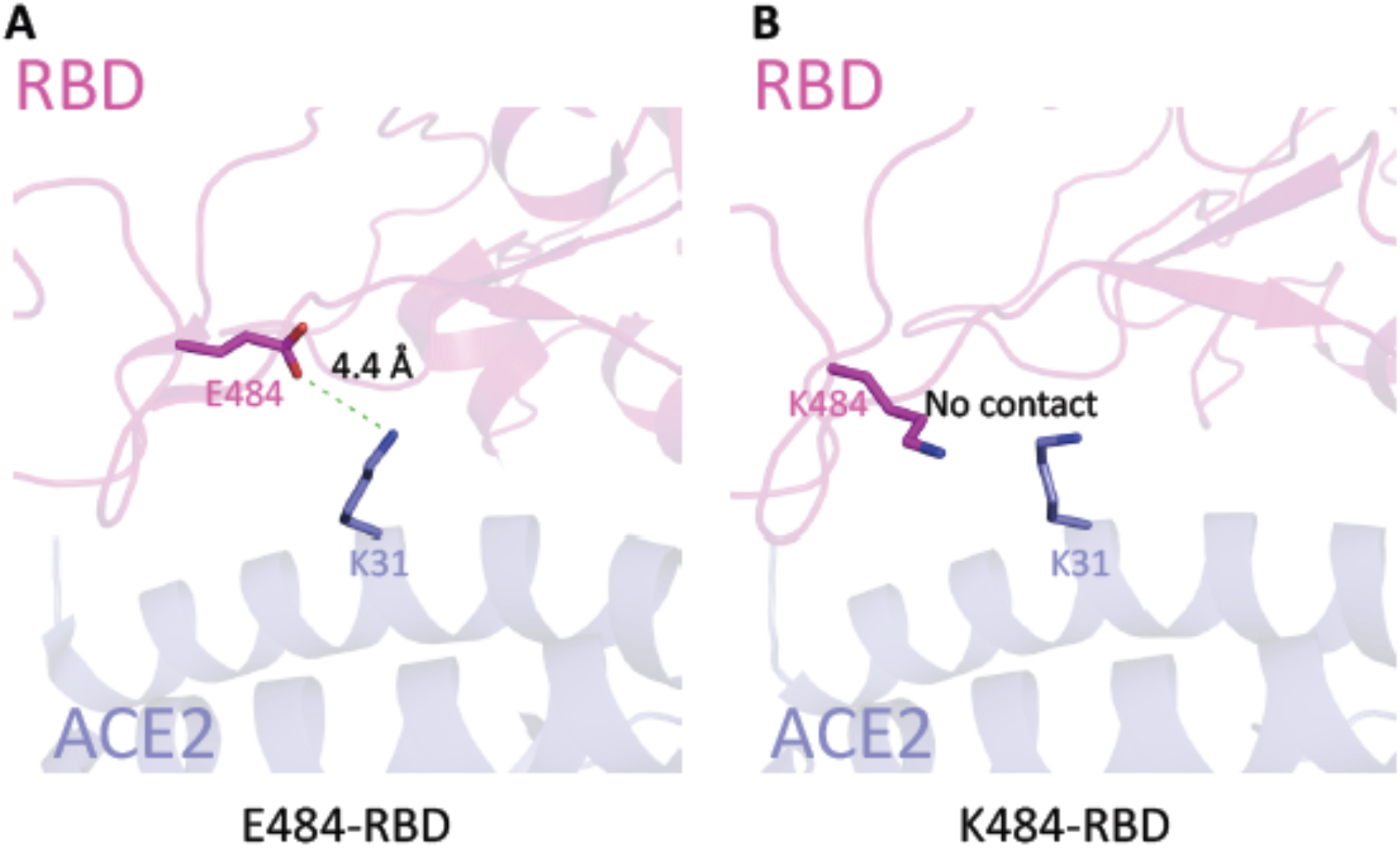
Model of E484K mutant. A. The interaction between E484-RBD and ACE2. B. The interaction between K4840RBD and ACE2.

Confirming our expectation, this additional mutation of K to N drastically reduced the binding affinity (~2.924 nM) between N417/Y501-RBD and ACE2 compared to 0.566 nM for the Y501-RBD to ACE2 interaction (**Fig. 1C**). Again, we did molecular a docking analysis of this mutation. The original salt bridge formed between K417 of RBD and D30 of ACE2 was lost in the new interaction but was replaced by a new weaker hydrogen bond (**Fig. 3A, 3B**). Overall, the changes of affinity with each mutation could be interpreted by the structural modeling data. We conclude that the three mutations of RBD from both the South African variant and the Brazilian variant lead to ~2 fold increase of binding affinity between the original RBD and ACE2, which may account for the higher infection rate of the South African and Brazilian variants over the original virus. owever, the affinity of these new variants is decreased by a factor of 5 compared to the Brisith variant. We demonstrated here how each individual muation contributes to the change of binding affinity either enhancing or reducing.

**Figure 3.**
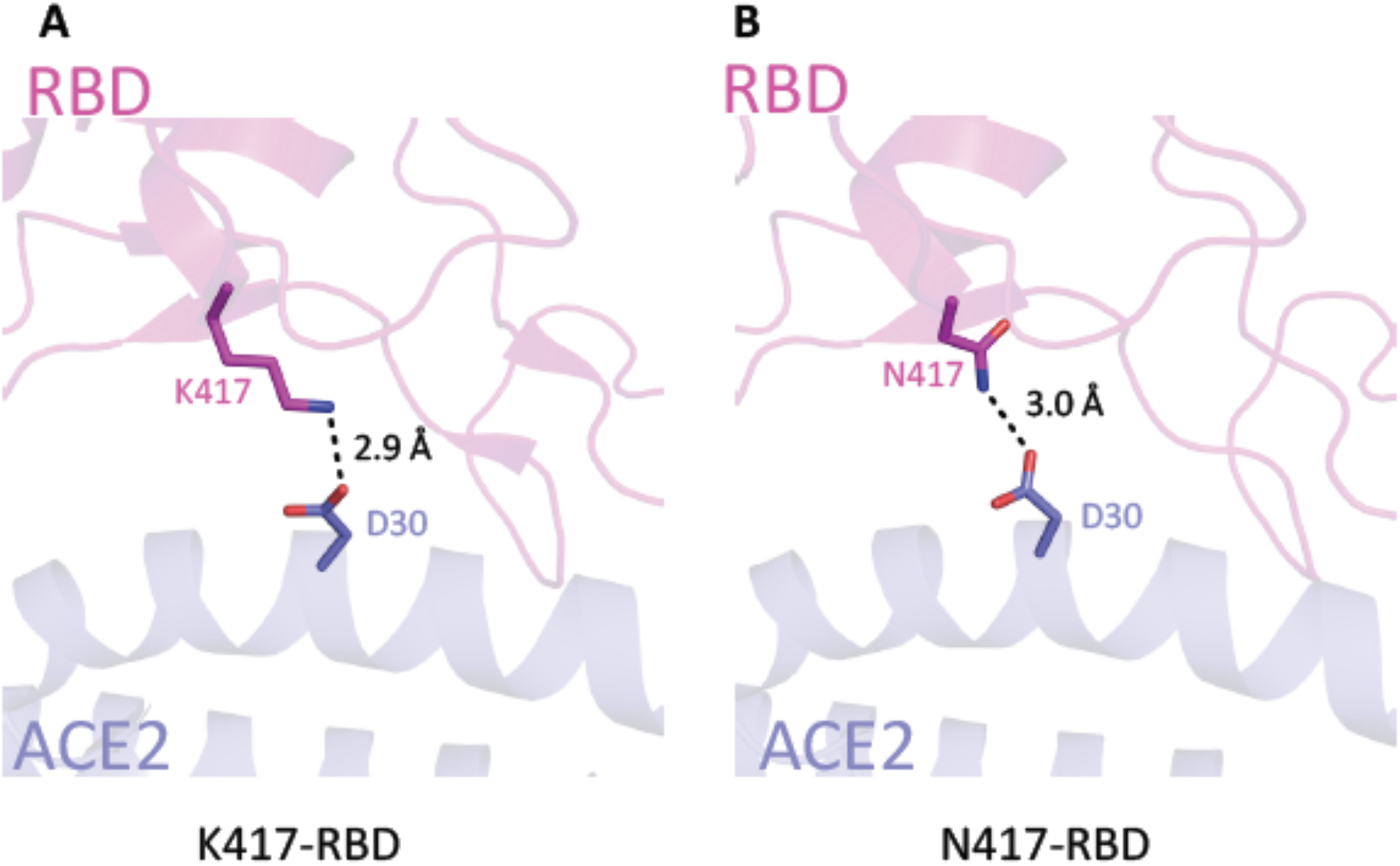
Model of K417N mutant. A. The interaction between K417-RBD and ACE2. B. The interaction between N417-RBD and ACE2.

Several monoclonal antibodies have been approved for therapeutic drugs to treat COVID-19 patients (Baum et al. 2020; Gottlieb et al. 2021; Bamlanivimab 2020; casirivimab and imdevimab 2020). Most importantly, all these therapeutic antibodies are RBD specific. It is of interest to investigate the binding of the mutant RBD with the therapeutic antibodies to verify the efficacy of these antibodies. The binding assays between three versions of RBD proteins with different mutation combination and Bamlanivimab were generated through applying them to immobilized antibodies on the Biacore Chip (**Fig. 4**). To our surprise, the interaction between the mutant RBD containing three mutations, N417.K484/Y501-RBD, and Bamlanivimab, was completely abolished (**Fig. 4A**). However, interestingly, the binding between N417/Y501-RBD and antibody remains the same as those between wildtype N501-RBD or mutated Y501-RBD and the antibody with binding affinities of 0.976 nM (**Fig. 4B**), 0.874 nM and 0.80 1 nM, respectively (Liu et al. 2021). Furthermore, as long as E484K mutation is introduced, the binding between K484/Y501-RBD or N417/K484/Y501-RBD and the antibody is completely abolished (**Fig. 4A, 4C**). The structure of the entire Spike protein and Bamlanivimab from Eli Lilly has been reported (Jones et al. 2020). We use this available complex structure to analyze how this critical E484K mutation causes this loss of binding. The structure showed that E484 of RBD forms three salt bridges with R50 and R96 of Bamlanivimab (LY-CoV555) (**Fig. 5A**). Our prediction shows that the conversion of E484 of RBD to K484 leads to the loss of three salt bridges as well as the formation of a new repulsive charge-charge interaction brought in from both positively charged sidechains of Lys and Args (**Fig. 5B**). This easily can explain the dramatic drop or loss of binding between RBD and the therapeutic antibody. From our binding data and structural analysis, we can conclude that Bamlanivimab (LY-CoV555) completely loses its efficacy in COVID-19 patients infected with the South African and Brazilian variants.

**Figure 4.**
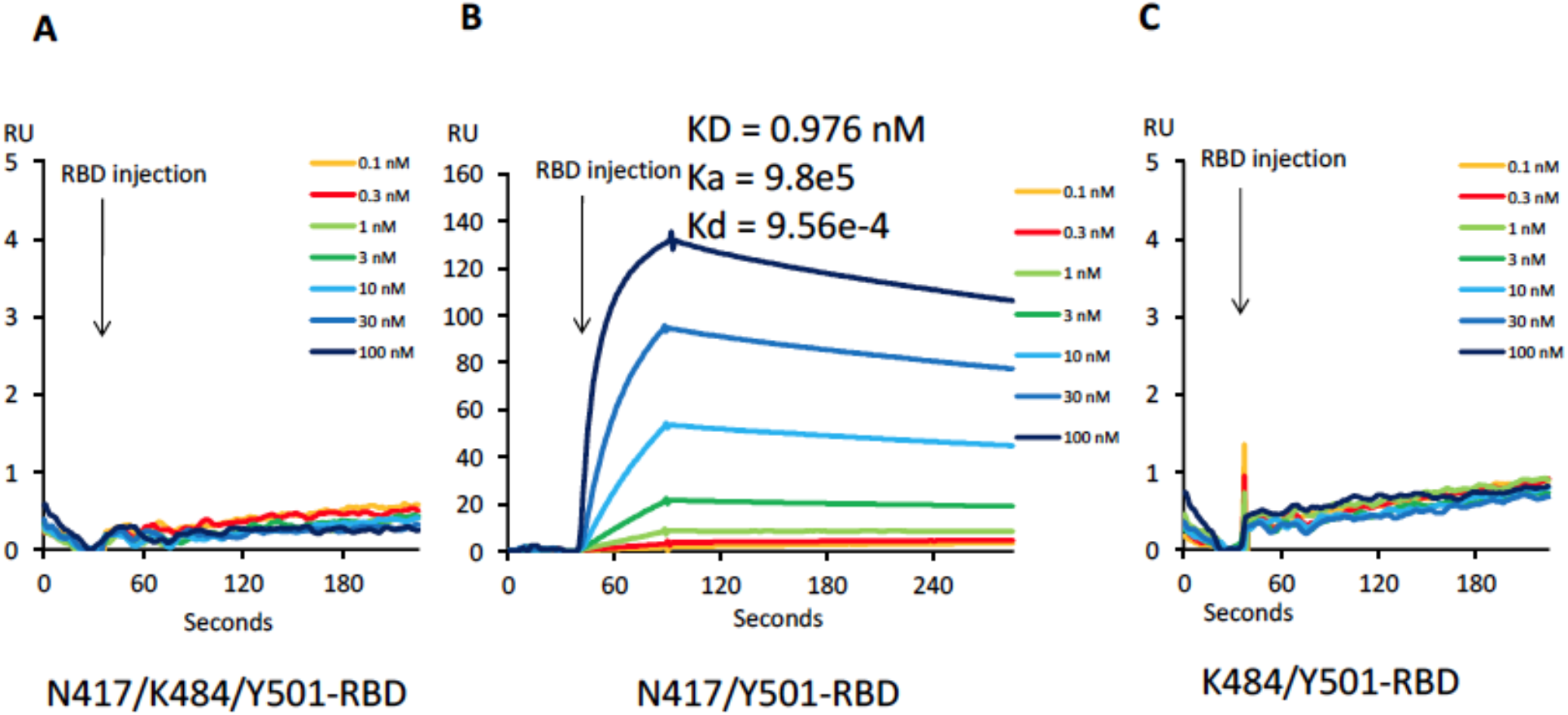
The basis of binding between mutated RBDs and the therapeutic antibody, Bamlanivimab. A. The binding between N417/K484/Y501-RBD and Bamlanivimab. B. The binding between N417/Y501-RBD and Bamlanivimab. C. The binding between K484/Y501-RBD and Bamlanivimab.

**Figure 5.**
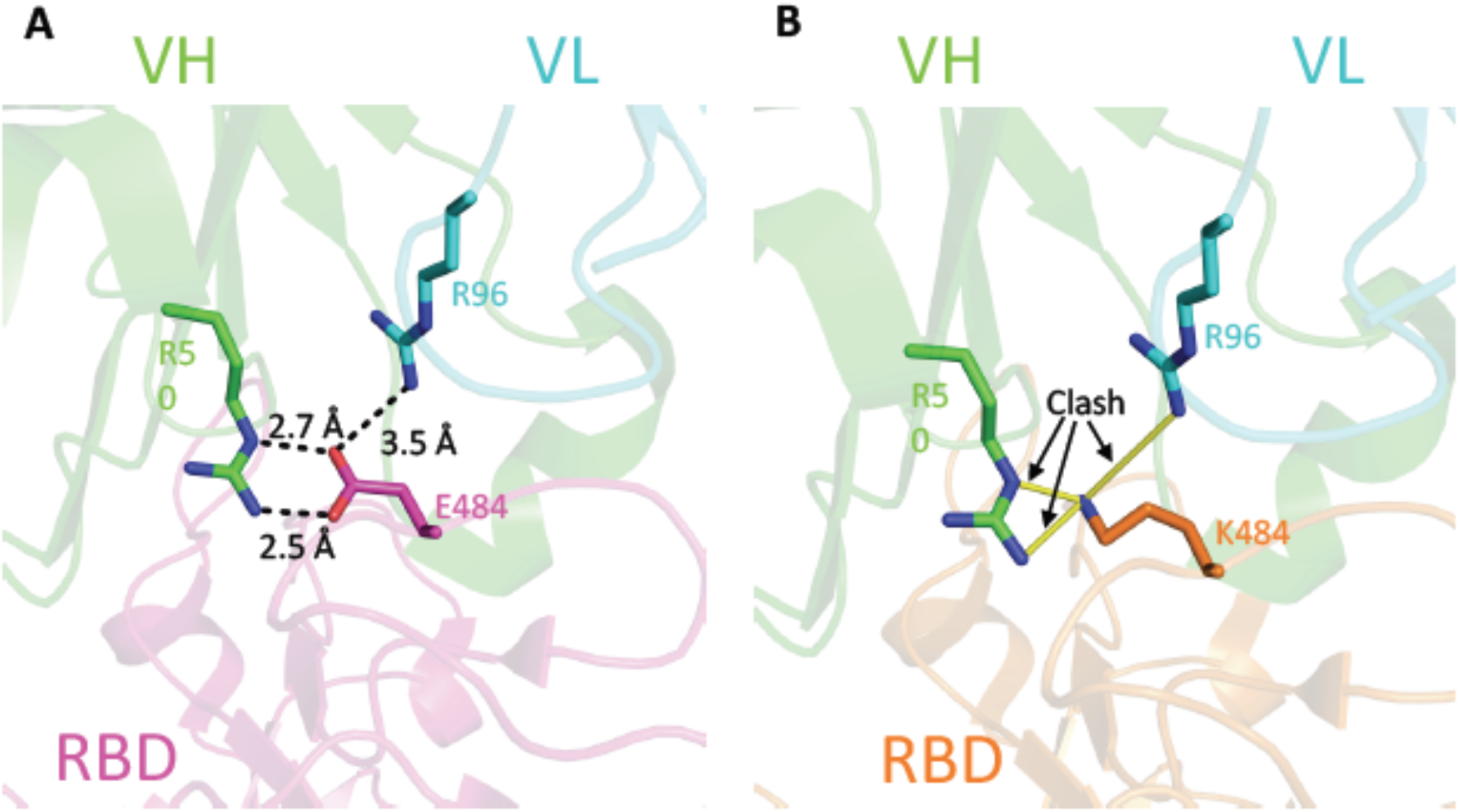
Model of interaction between mutant RBD and Bamlanivimab. A. The interaction between E484-RBD and Bamlanivimab. B. The interaction between K484-RBD and Bamlanivimab.

The COVID-19 pandemic has caused disasters for humans, and it is still out of control with limited means to contain it. Any mutation associated with an increased rate of infection could be a warning sign of potential escape of the virus from existing therapeutic options and pose challenges to both the available vaccines and therapeutic antibodies. Luckily, the mutation rate of the virus is low. Nevertheless, mutations in critical sites still could threaten the efficacy of vaccines and therapeutic neutralizing antibodies. At the moment, researchers currently pay mostly attention to the efficacy of broadly neutralizing antibodies generated by COVID-19 patients and vaccines. As explicitly demonstrated by Dr. Shane Crotty and his colleagues (Dan et al. 2021; Grifoni et al. 2020; Rydyznski Moderbacher et al. 2020; Sette and Crotty 2021), the T cell arm of adaptive immunity actually plays a similar critical if not the more important role in battling the virus. While B cells use strictly 3D epitopes from the virus, helper T cells and cytoxic T cells utilize linear epitopes in their fight against the virus. Thus, even with the loss of efficacy of broadly neutralizing antibodies against the RBD regions of occasional escaping variants, T cells with their completely different protection mechanism through conserved linear antigens derived from SARS-CoV-2 could build up a strong defense firewall against newly emerging and slowly evolved SARS-CoV-2 variants making the loss of efficacy of antibodies a less frightening prospect. We believe that broad vaccinations with two mRNA based vaccines from both Pfizer-BioNTech and Moderna, or others will build up herd immunity to beat SARS-CoV-2. However, vaccines that provide a large group of T cell antigens of the entire virus, rather than just the Spike protein, for example inactivated virus vaccines, should not be forgotten. This report shows the danger of mutations within the surface of the ACE2 binding region of RBD from the aspect of therapeutic antibody efficacy. These therapeutic antibodies were derived from a few conserved 3D epitopes of the virus, especially the RBD domain. We showed previously that with respect to the United Kingdom variant, it may require much higher concentration of the antibody to treat COVID-19 patients with the United Kingdom variant compared to patients with wildtype virus, even though it does not affect the binding of therapeutic antibody Bamlanivimab (Liu et al., 2021). On the other hand, some additional mutations within the RBD regions, such as E484K besides N501Y within both the South Africa and Brazilian variants, cause complete loss of efficacy of Bamlanivimab. To be clear, this is *in vitro* data, it needs further verification *in vivo*. From this perspective, one needs to be aware that therapeutic antibodies to treat COVID-19 patients should be adjusted for new emerging variants accordingly.

## Methods

### SARS-CoV-2 spike RBD mutation and expression

SARS-CoV-2 RBD (319-541aa) was cloned to pCDNA3.1 vector with 6 histidine tag at the C-terminal. Y501 mutation was created by quick change mutagenesis and verified by DNA sequencing. N417 or K484 was mutated on the Y501-RBD backbone by the same method to create RBD with double mutation sites. The triple mutation RBD was created in the same manner. All the mutations were verified by DNA sequencing. The mutated RBD was transiently expressed in 293F cell line by transfection. The supernatant was used for RBD purification by passing through the nickel column. The eluted RBD was further purified by Superdex-200 Gel-filtration size column.

### Biacore affinity measurement of mutant RBD binding to ACE2

Affinity measurement was carried out with Biacore T200. Biotinylated BirA tag-ACE2 (1-615) was used to coat Strepavidin chip (Liu et al. 2021). Mutant RBD was injected at different concentrations. The affinity was calculated by the BIA evaluation software.

### Biacore affinity measurement of mutant RBD binding to Bamlanivimab

Affinity measurement was carried out with Biacore T200. Bamlanivimab was coated on the CM5 chip by the amine coupling method (Liu et al. 2021). Mutant RBD was injected at different concentrations. The bound RBD was eluted with 10 mM glycine pH 1.7. The affinity was calculated by the BIA evaluation software.

### Protein docking and binding affinity prediction for mutants

Protein docking studies were performed by HADDock 2.2 server(Van Zundert et al. 2016). Crystal structure of SARS-CoV-2 RBD bound with ACE2 (PDB code: 6M0J) was selected for docking. Two different mutants (N501Y_K417N and N501Y_E484) structure was prepared by Coot software(Emsley and Cowtan 2004). One of the conformations from alternative residues in the structure was deleted manually to meet the docking server criteria. The major contacts residues in 6M0J (hydrogen bond and salt bridge) were selected as restrain residues for docking (Q24, E35, E37, D38, Y41, Q42 and Y83 for ACE2 protein; N487, Q493, Y449, and Y505 for RBD protein). The best model from the docking was selected for the next analysis. All protein structural figure prepared by PyMOL. The binding affinity prediction for each of mutants was did by BeAtMuSiC server(Dehouck et al. 2013).

## Acknowledgements

We thank National Jewish Health for supports and people in the Kappler/Marrack groups for help. H.L. is partially supported by NIH grant (GM135421 to G.Z,) and NB Life Laboratory LLC. We also thank the Colorado Department of Public Health and Environment (CDPHE) for authorization to use the residual therapeutic antibodies after COVID-19 patient infusions.

## Contributions

H.L., P.M., and G.Z. for designing; H.L. for main experiments; P.W., Q.Z., Z.C., K.A., W.D., S.P., G.D., L.R., S.F. for some experiments; H.L., P.M., G.Z. for final data analysis and writing up.

